# Polyol sugar osmolyte - sorbitol corrects chromosome cohesion and translational defects in cohesin mutants

**DOI:** 10.1101/2024.09.04.610945

**Authors:** Sayali Marathe, Haripriya Chougule, Vandana Nikam, Amitabha Majumdar, Tania Bose

## Abstract

The cohesin protein complex plays a very important role in chromosome segregation, transcription, DNA replication and chromosome condensation. Mutations in cohesin proteins give rise to a disease collectively referred to as Cohesinopathies. The major cause of Cohesinopathies arise due to defects associated with gene expression, that give rise to developmental disorders. In this study, we have used *Saccharomyces cerevisiae* to mimic the Cohesinopathy disorder Roberts syndrome with mutations (*eco1W216G*) homologous to that of humans (esco2). Our data suggests that polyol sugars like sorbitol, can repair misfolded proteins and reduce ER and proteostatic stress. We have used sorbitol as a chemical chaperone, to check how it can restore chromosome segregation, gene expression, misregulation, protein misfolding, autophagy and translational defects in the cohesin mutant of the Roberts’ phenotype. *In silico* screening has helped us identify the possible sites on eco1, which could be possibly altering the phenotypic traits.

**Article Highlights:** *In presence of sorbitol:*

- **Temperature sensitivity of the Roberts mutants is rescued.**
- **Chromosome cohesion defect is reduced.**
- **Translational defects are minimized.**
- **Autophagy is enhanced**
- **Molecular docking shows its interaction with the acetyltransferase domain of ESCO2.**

## 1. Introduction

Cohesin complex is composed of a group of proteins such as SMC1A, SMC3, and RAD21 (MCD1 in yeast). These proteins are loaded onto chromosomes by NIPBL^1^. Cohesin proteins bind to the sister chromatids and maintain cohesion between chromatids along the entire length of chromosome ^2^. These remain attached till metaphase till proper separation of sister chromatids occurs at anaphase ^2^. In addition to the cohesin complex, there are acessory proteins which play an important role in providing stability, and maintaining genomic integrity ^3^ such as eco1, pds5 etc. These proteins are cell cycle-dependent. Mutations in any of these proteins, lead to diseased conditions referred to as Cohesinopathies ^3^. There are different types of cohesinopathy disorders caused by mutations in RAD21, NIPBL, SMC1A, SMC3 and HDAC8 which show some common phenotypic characteristics. Amongst all these syndromes, Cornelia de Lange Syndrome (CdLS) and Roberts’ are well explained. CdLS is caused by a mutation (heterozygous) in nipbl, smc1a and smc3 ^1^ whereas mutation in *eco1-ack, eco1-1* and *eco1W*216*G* (esco2), causes Roberts’ syndrome ^4,5^. These mutations affect growth and delays development.

In this study, we have studied the effects of Roberts’ syndrome (RS), which is a rare autosomal genetic disorde. The disease manifestations in RS are stronger than CdLS. Roberts syndrome is characterized by limb and craniofacial defects. Cases have been reported with abnormalities in the heart, urinary tract and other organs, in addition to stillbirth and sudden mortality in the early stages of development ^6^. It is caused by a point mutation in the esco2 gene (yeast eco1) ^4,7^, resulting in loss of function. In Roberts’ mutation, heterochromatic regions show precocious separation of sister chromatids, which lead to development of micronuclei and aneuploidy ^8^.

It is important to understand the pathophysiological effects caused due to these mutations. Different models such as yeast, mice, zebrafish and *Drosophila* have been used to study the disease mechanism of RS ^9^. For our work we have used the haploid yeast model, to introduce mutations in eco1, analogous to the human mutation ^7^. The Roberts’ phenotype shows cohesion defects, which can be detected by chromosome cohesion assays ^10^. Also, this mutant shows moderate temperature sensitivity at 33°C, lethality at 37°C and thrives best at an optimum temperature of 30°C. The nucleolar morphology, adjacent rDNA disruption, cohesin less binding of cohesin rings and translation by RNA Pol I is affected ^11^. In addition, to chromosome segregation, there are also defects at the transcriptional level which give rise to developmental disorders ^3,4^.

The RS mutant is a stress response mutant, associated with defects in protein translation ^4,12^. From our previous work and those of other groups, we know that cohesin proteins are involved in the regulation of translation ^4,13^. Our study aims to investigate how the chemical chaperone sorbitol, can alter the elevated expression of the stress response genes in the mutant as well as correct chromosome segregation, translation, protein misfolding and autophagy. We have also attempted to check for possible interactions with sorbitol, by using *in silico* approach.

Due to translational defects, there is an accumulation of misfolded proteins ^13^. These misfolded proteins tend to aggregate in the endoplasmic reticulum (ER) ^14^, generating cellular stress ^15^. To rescue cells from stress there is either a shutdown of transcription or activation of various pathways like unfolded protein response (UPR), stress regulation and CWI (Cell Wall Integrity) pathway ^16^. Due to changes in expression levels of all these proteins, newly synthesized misfolded proteins, are minimally produced, which gives more time for degradation pathways to dispose of unwanted residue ^17,18^. Moreover, changes in gene expression of protective sugars help the cells to survive stressed conditions ^18^. Similar to neurodegenerative diseases ^17^, it is hypothesized, in the *eco1* mutant, there is a built up of ER stress which causes accumulation of unwanted proteins. Under these conditions, it tends to increase the expression of chaperones ^19^, to degrade abnormal and toxic proteins which might correct the misfolding. This helps, to rescue the cells from the stressed situation ^20^.The proteostatic stress in the cells is measured by luciferase assay, where sensor proteins from firefly are used. These sensor proteins tend to misfold during stressed condition. After misfolding it loses its luminescence activity ^21^.

These misfolded proteins are corrected with the help of molecular chaperones ^20^, but due to continuous production of misfolded proteins, it increases the load on the chaperone machinery. Researchers have used external chaperones, which assist misfolded proteins to fold properly and regain their function ^10, 20–22^. The use of cellular chaperones is known for correcting misfolded proteins and ultimately reducing the severity, observed in case of many of the neurodegenerative diseases ^24^. It has been reported, that chemical chaperones are used in the treatment of various neurodegenerative diseases, like Huntington, Amyotrophic lateral sclerosis, Alzheimer and Parkinson’s disease ^25,26^.

Numerous reports suggest use of low molecular weight compounds such as glycerol, TMAO ^27^ and other amino acid derivatives, naturally present in cells at higher concentrations, referred to as osmolytes or chemical chaperones. These chemical chaperones can reduce and reverse the aggregation of misfolded proteins ^25^. They play an important role in providing stability to proteins, due to its ubiquitous action. These are well known for stabilizing the proteome under natural conditions and reduce ER stress ^28^. It can be widely used by various organisms for intrinsic cellular processes. Hence chemical chaperones can be considered as a potential alternative to biological chaperones ^29^. The exact mechanism of action of these chemical chaperones is not clear, but it probably functions by preventing nonproductive interactions as well as by altering endogenous activities of the chaperones. This stabilizes misfolded proteins and ultimately reduces protein aggregation ^25^.

The chemical chaperones are either polyhydric alcohol and sugars (like trehalose, sorbitol, glycerol, arabitol, mannitol and myo-inositol) ^26^, amino acids derivatives (Proline, Glycine, Alanine, Taurine) or methyl ammonium derivatives (Trimethylamine N-Oxide i.e. TMAO, Glycreo-Phosporyl choline). Hence depending on the mechanism of action for generating stability, these osmolytes are classified either as compatible or counteracting osmolytes. The first class of osmolytes are compatible type of osmolytes which include aminoacids and polyols like sorbitol. These do not show marked effect on the functional properties of the protein and provide stability under native conditions. The second class is the counteracting class, which causes modification in the functions of proteins, helps to provide stability and inactivates the protein. These include methyl ammonium derivatives ^30,31^.

Proteins, in the presence of osmolytes, are far more stable than those in the absence of the osmolytes under denaturing conditions. These osmolytes enhance protein folding and stability of the protein ^32^.

During protein folding, osmolytes help in stabilizing proteins by interacting with the peptide backbone of the protein. This interaction allows protein domains to gain their preferred hydration ^32^. Compounds like sorbitol tend to avoid direct contact with polypeptides. Hence, at higher concentrations, we observe increased hydration around polypeptides, as proteins pack themselves more tightly and decrease their surface area. This phenomenon of packing of polypeptides in the presence of osmolytes increases the stability of the proteins ^30, 31^. It has been reported that the presence of these polyols, glycine and betaine are some of the best osmoprotectants which preferentially exclude water molecules and increase the thermal stability of proteins ^33^. *In-vitro* and *in-vivo* studies have shown that amyloid aggregation of misfolded proteins with a fibrillar nature, are deposited in cells during ER stress, get reduced in the presence of polyols ^29^.

Sorbitol (glucitol) is a sugar alcohol/polyol and falls under the group of compatible type of osmolytes. It is widely used as a stabilizing agent in food and pharmaceutical industries ^33^. Sorbitol remains stable at all temperatures and is a chemically inert molecule which does not have significant effect on the functional properties of proteins ^31^. It helps to provide stability to proteins and maintain their native conformation ^30^. Other than sorbitol, mannitol (which is also a 6-carbon sugar differs in positioning of the -OH group of sorbitol), is effective as a chemical chaperone. Mannitol can temporarily disrupt the blood-brain barrier and inhibit aggregation. Hence, it can be used as a drug in the treatment of Parkinson’s disease. Mannitol also stabilizes proteins by increasing its concentration inside the cell through the process of osmosis. It also shows neuroprotective effects on transgenic mice ^34,35^.

In this study, we have chosen sorbitol as a chemical chaperone ^36,37^, to check, if it can rescue the diseased condition in the Roberts’ mutant. It has been reported that it can pass through the blood-brain barrier, has a short half-life in serum, and hence will not accumulate in blood. Sorbitol is easily metabolized by the brain. It can be used as an effective osmotherapeutic drug at optimal doses ^38^. Taking into view, the following characteristics of sorbitol, we have used it as a chemical chaperone which can possibly rescue the diseased condition of RS in yeasts. It has been reported that due to hyperosmolarity of sorbitol, there is a change in the transcription and translational processes, cell cycle progression and retention of various osmolytes ^39^.

In untreated samples, wild type and mutant cells are lethal at 37°C, shows slow growth at 33°C and optimum growth at 30°C. These mutants show chromosome cohesion defects, levels of phosphorylation of eif2α is enhanced, autophagy is reduced, in comparison to wild type cells.The defects in translation initiation and misfolded proteins accumulated in ER is also observed in untreated mutant cells.

In this article we report that post-treatment of wild type and mutant cells with 0.5M, 1M, 1.5M, and 2M concentration of sorbitol, the results from the growth and chromosome cohesion (one spot two spot) assay showed rescue of the mutant at 37°C. Phosphorylation of eif2α is reduced and autophagy levels are enhanced in the mutant, actively translating ribosomes increase in sorbitol-treated samples which are almost at the same level observed under normal conditions. The results obtained suggest that sorbitol possibly acts as a rescuer of the phenotypic manifestations of the diseased condition observed in RS. Altered autophagy is one of the leading causes of protein misfolding. Under these conditions, there is a built up of protein in the system. Reduction in autophagy, shifts the flux of misfolded proteins resulting in proteotoxicity ^40^. Cells can function continuously to remove these misfolded proteins; however, if this clearance mechanism including autophagy is affected, it will lead to built up of toxic proteins within cells leading to lethality ^41^. Molecular docking studies showed that sorbitol possibly binds to the acetyltransferase domain of human esco2, altering the deleterious effects of the mutation.

## 2. Materials and methods

### 2.1. Serial dilution assay

Wild type and *eco1*W216*G* strains were grown in YPD broth overnight. The next day O.D. (Optical Density) was adjusted to 0.2. The cultures were serially diluted and spotted on YPD plates with different concentrations of sorbitol (0.5M, 1M, 1.5M, 2M) and incubated for 48 hours. These growth plates were studied in 3 biological replicates. After 48 hours results were checked for growth pattern and further analyzed ^42^.

### 2.2. Growth Assay

Wild type and *eco1W216G* cultures of yeasts were grown in YPD broth till it reaches mid-log phase. Then cultures were grown in a 96-well plate containing YPD with sorbitol and YPD medium as control. This was performed in three biological replicates. Each batch was set up for 48 hours at 30°C with continuous shaking in the Epoch2 BioTek Spectrophotometer and O.D. was recorded every 30 min. The data was acquired and the graph was plotted using GraphPad Prism 9 software ^43^.

### 2.3. Microscopy

To check chromosome cohesion by microscopy wild type and *eco1W*216*G* strains of yeast with 50 and 100 copies of rDNA were grown overnight in YPD medium. The cells were arrested with arrested and stained with Hoechst staining^7^. Culture was placed on a slide and observed under a Nikon Ti Eclipse microscope with GFP laser using 60x and 100x objectives for presence of one or two GFP spots. All the images were acquired at 60X zoom using a 100X plan objective, using the manufacturer’s software. FIJI software was used to process Images. The cells showing one spot and two spots were counted and 100 cells from different fields were calculated. Graphs were prepared using GraphPad Prism 9 software ^4^.

### 2.4(a). RNA sequencing

Wild type and *eco1W*216*G* strains were grown in YPD with and without sorbitol in a shaking incubator overnight. The RNA was isolated using the Ambion RNA extraction kit. The sample was quantified and RNA sequencing was performed using Illumina sequencing method.

The threshold for categorizing the data into up and down is +/-1 where > = +1 log2Fold change are upregulated genes < = −1 are downregulated and between +1 and −1 are neutrally regulated. In order to generate the heatmap, we have sorted the ‘log2fold change’ column "B" in descending order and plotted the top 20 up and downregulated genes along with the normalized values.

The colour key represents the regulation where the green colour represents positively regulated genes or an increase in expression and the yellow colour represents downregulated genes or a decrease in expression.

### 2.4(b). RT- PCR

Cultures of wild type and *eco1W*216*G* were grown in YPD with and without sorbitol, overnight, in a shaking incubator. The RNA was isolated using the Ambion RNA extraction kit method. The RNA was treated with Turbo DNase. The RNA (2ug) was then quantitated using nanodrop. cDNA was made using a dNTP mix and Random 6 mers and Takara cDNA kit. RT- PCR was performed using cDNA (5-fold dilution), 18S and 25S forward and reverse primers. We have used actin as a housekeeping gene. Gene expression values were plotted using GraphPad9 software ^4^.

### 2.5 Luciferase sensitivity assay

Strains of wild type eco*1W*216*G* harbouring the FlucDM plasmid were grown at 30 °C till logarithmic phase in 5ml of media containing yeast extract, peptone and 2% dextrose. Culture was spin down at 3000xg or 10 min. Pellet was washed with water and resuspended in lysis buffer (Tris 50mM, Sodium chloride 150mM, NP-40 0.1%, Dithiothreitol 1mM,Glycerol 10% and Protease Inhibitor).Cells were added with 3-4 beads and lysis was done with bead beater. Lysate was collected and centrifuged for 10,000 rpm for 15 min at 4 °C. Luminiscence was assesed by adding 100µL lysate to equal amount of Firefly Luciferase Assay Substrate (Promega).The mixture was incubated for 5 min and reading was taken on a luminometer. Data was analysed with Grahpad Prism 9 software ^21^.

### 2.6. Western blot

Cultures of wild type and *eco1W*216*G* cells (in presence and absence of sorbitol with YPD as a the medium) were grown to an OD_600nm ∼_ 0.8. The cells were pelleted by centrifugation followed by lysis using lysing matrix containing glass beads and lysis buffer (Tris10 mM (pH 7.4), 1mM Sodium Chloride, 1 mM EDTA100 mM, EGTA 1 mM, Sodium Fluoride 1 mM, Tetrasodium Pyrophosphate 20 mM, Sodium Orthovanadate 2 mM, Sodium Dodecyl Sulfate 0.1%, sodium deoxycholate 0.5%, Triton-X 100 1%, glycerol10%, and PMSF1 mM, protease inhibitor cocktail (Sigma)). The cell lysis was done using a bead beater (MP biomedicals^TM^-116005500)followed by centrifugation. The supernatant was collected and quantified using the Bradford method. Equal amounts of protein were loaded for every sample. Protein samples were separated by SDS gel electrophoresis using 12% SDS-polyacrylamide followed by its transfer on nitrocellulose filters. After transfer the blots were probed with primary antibody phospho-specific eif2α (Ser-51) (Cell Signaling). This is followed by a secondary antibody (anti-rabbit). The bands were visualized by using developer(Thermo) and Amersham imager 600 (GE Healthcare Life Sciences) ^4,44^.

A similar procedure of western blotting was followed to check autophagy levels. The detection was done using ATG8 (Thermo Fisher Scientific) as primary antibody ^45^ and anti-Rabbit antibody as secondary antibody. Bands were visualized using developer (Thermo Fisher Scientific) and Amersham imager 600 (GE Healthcare Life Sciences).

### 2.7 Polysome profiling

For polysome analysis 200 ml of yeast culture was grown up to an OD_600nm_ 0.8. Cells were pelleted down washed with PBS and treated with 20 mg/ml of cycloheximide for 10 mins on ice. After 10 min, cells were subjected to centrifugation. The obtained cell pellets were added in lysing matrix tubes containing glass beads and lysis buffer (Tris-HCl10 mM pH 7.5, Sodium Chloride100 mM, Magnesium Chloride 30 mM, Cycloheximide100 ug/ml, heparin 0.2 mg/ml in DEPC). This is followed by cell lysis using cells bead beater (MP Biomedicals) centrifuged and 500µl of supernatant was loaded on top of an 11 ml 7–47% sucrose gradient prepared in High Salt Resolving Buffer (Tris-Cl 15 mM pH 7.4, Ammonium Chloride 140 mM and Magnesium acetate tetrahydrate 7.8 mM). Tubes were then subjected to ultracentrifugation at 27,000 rpm for 2.5 h in swing bucket SW41 rotor. After centrifugation, the gradients were fractionated, and OD was monitored at 254 nm using a Bio comp fractionator station. Peaks were analyzed after measuring polysome to monosome ratio ^46,47^.

### 2.8 In silico screening

The preparation of target proteins, ligand and molecular docking were done as described earlier ^48^. Briefly, the target proteins, Esco2 acetyltransferase in complex with CoA (PDB ID – 6SP0) was downloaded from RCSB Protein Data Bank with 1.77Å resolution. The target protein has two ligands Zn and Coenzyme A (CoA).

The target proteins in protein data bank (pdb) format were saved and protein preparation was done using the AutoDock Tools (version 1.5.7). The output grid dimension files were saved after adjusting the grid on the CoA site of the target proteins. The grid dimensions at CoA site with ligand Zn without CoA were center_x = 20.483;center_y = 46.761; center_z = 67.197 for CoA site. The blind docking was performed with keeping both ligands, Zn and CoA, intact with grid dimension center_x = 21.975; center_y = 39.649; center_z = 64.832

The ligand structures were downloaded from PubChem database, energy minimized using Chimera (version 1.15) and preparation was done using AutoDock Tools (version 1.5.7). Molecular docking was done using AutoDockVina ^48^.

The protein-ligand interactions were visualized using Biovia Discovery Studio 2021 and validation of molecular docking protocol (Root Mean Square Deviation - RMSD) was done using PyMOL software.

## 3. Results

### 3.1. Temperature sensitivity of eco1 mutant strain rescued in presence of sorbitol

The wild type and mutant cells were checked for 48 hrs in the presence of different concentrations of sorbitol (0.5M, 1M, 1.5M, 2M) by spot dilution assay. Due to mutation and translational defects, these cells show moderate growth at 30°C, slow growth at 33°C **(Fig 1)** and lethality at 37°C. After 48 hrs we observed that YPD plates that had been incubated at 37°C show no growth at 0.5M and 1M, whereas in presence of 1.5M and 2M sorbitol, the growth was revived.

**Fig.1(a).**
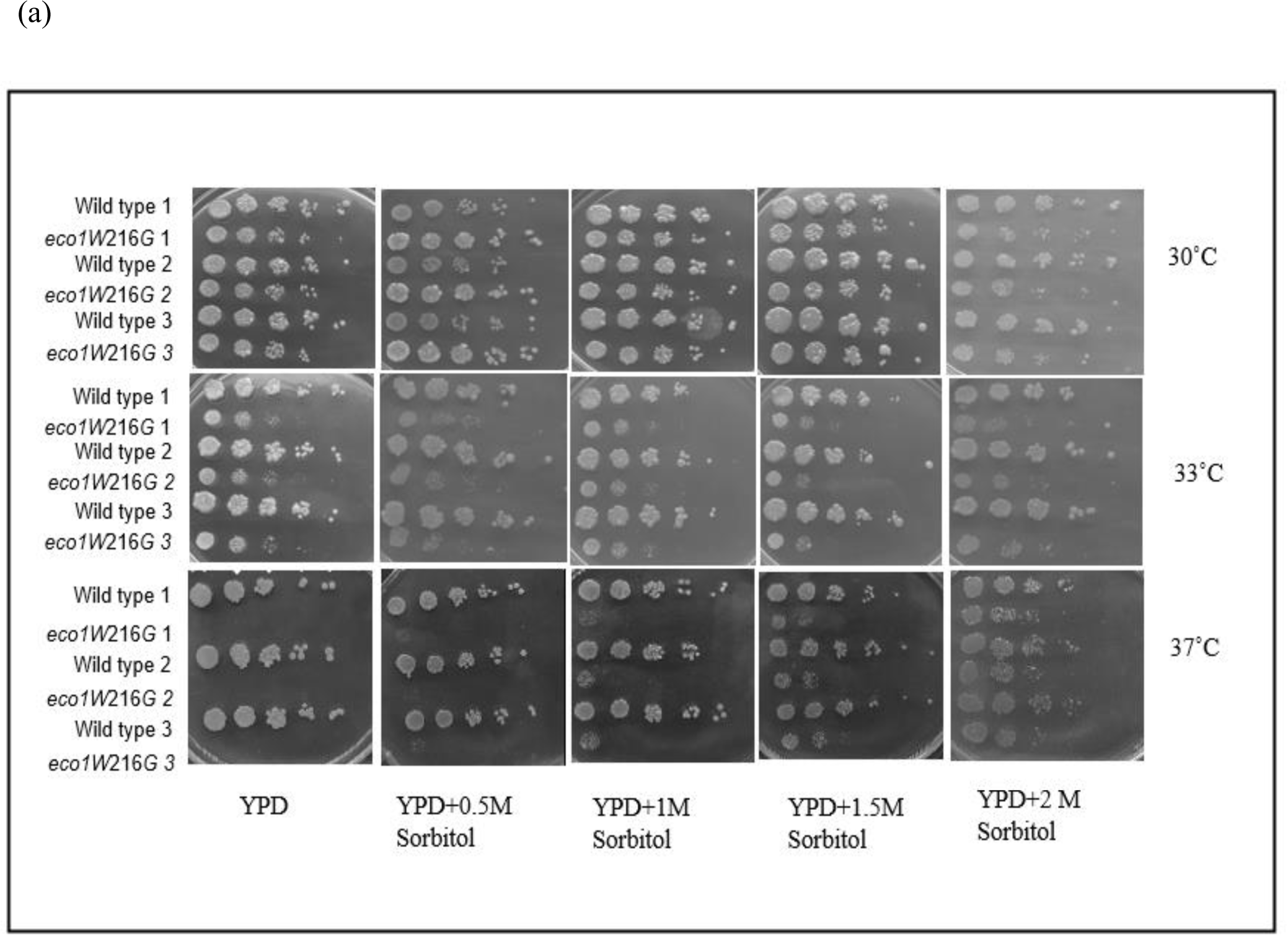
Effect of sorbitol on the growth of wild type and mutant (*eco1W216G*) at 37°C (serial dilution assay) Wild type and mutant (*eco1W*216*G*) strains of *Saccharomyces cerevisiae* were grown, and serially diluted and spotted onto YPD medium containing 0.5M, 1M, 1.5M and 2M sorbitol respectively. The data shown here is a representative of three independent biological replicates.

**Fig.1(b).**
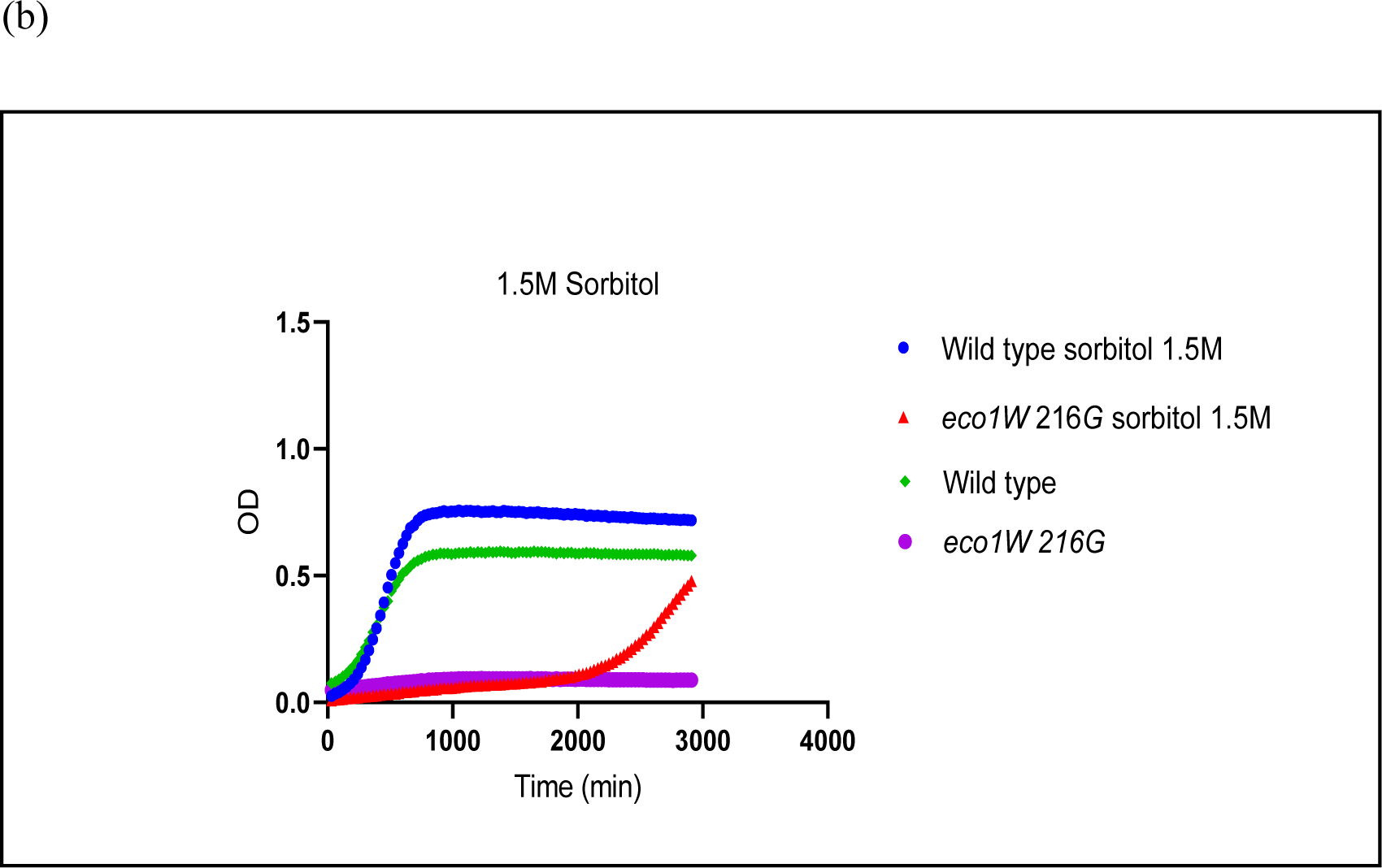
Effect of sorbitol on the growth of wild type and *eco1W216G* at 37°C (Growth Curve Assay) 48hrs growth curve of wild type and *eco1W216G* treated with 1.5 M with and without Sorbitol. The O.D._600_ was measured at 30 min intervals. Data shown is obtained from 3 independent biological replicates. Green line 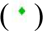 - wild type, Purple line 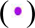 - *eco1W216G,* blue line 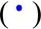 - wild type with 1.5 M sorbitol, red line 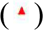 - *eco1W216G* treated with 2M Sorbitol, Graphs for growth curve were plotted using GraphPad prism 9 software.

We observed, in plates with 2M sorbitol, growth is revived, but there was a slight reduction in the growth of wild type cells at 37°C. Thus, we have selected 1.5M sorbitol concentration for further assays. After primary screening, we can hypothesize that sorbitol acts as a suppressor and the mutant shows a rescue of growth at 37°C.

To further consolidate the above results, we have analyzed the growth pattern of wild type and Roberts’ mutant cells at 37°C, using growth curve assay, where wild type and mutant cells were grown in YPD medium in the absence and presence of 1.5M sorbitol. We monitored the growth by following the absorbance at 600 nm at intervals of 30 min for 24 hrs. This data was analyzed using GraphPad Prism 9. We observed rapid growth with a very short lag phase in wild type and mutant cells inoculated in YPD in absence of sorbitol **(Fig 1)**. On the other hand, cells inoculated at 1.5 M took a longer lag phase but showed better growth following log phase, indicating rescue of the mutant cells in presence of 1.5M sorbitol.

### 3.2. Chromosome cohesion (one spot vs two spot) assay to check if sorbitol changes chromosome cohesion defects

Mutations in the acetyltransferase domain of *eco1w216G*, gives rise to gene misregulation. These are associated with changes in the nucleolar size. rDNA is an actively transcribed region situated adjacent to the nucleolus. This leads to changes in the morphology of rDNA as well ^11,44^. rDNA is also involved in the maintenance of ribosome biogenesis. Previous reports suggest, that, the Roberts’ mutant shows aberrant rDNA with a reduction in the number of ribosomes actively involved in translation ^11^. We have used wild type and mutant strains for checking chromosome cohesion, with 50 and 100 copies of the rDNA cluster respectively. An increased number of rDNA can cause a delay in replication and separation at anaphase. Variation in copy number of rDNA helps to detect chromosome cohesion in the presence and absence of sorbitol. The plasmid used for detecting chrosomosome cohesiveneness, consists of an array of lac repressor sites integrated into the chromosome IV- R arm with LacI-GFP system ^9^. To check if addition of sorbitol has any effect on chromosome cohesion we have used wild type and eco1 mutant, which were treated with nocodazole for arresting at G2/M phase. It was checked for the presence of one or two spots. Two distinct spots at one locus indicate precocious separation of sister chromatids before anaphase ^9^ and a single GFP spot indicates intact chromosome cohesion. Around 100 cells were checked and the data was analyzed using GraphPad Prism 9. The data suggests, that in untreated mutant samples, there were twice the number of GFP spots in comparison to the wild type cells, indicating more of the two spots. After treatment with 1.5M sorbitol, the number of two spots were reduced, indicating chromosome cohesion was restored in presence of sorbitol **(Fig 1)**. With both 50 and 100 rDNA copies we observed a similar pattern in the reduction of chromosome cohesion, which is corrected in presence of sorbitol.

**Fig.2.**
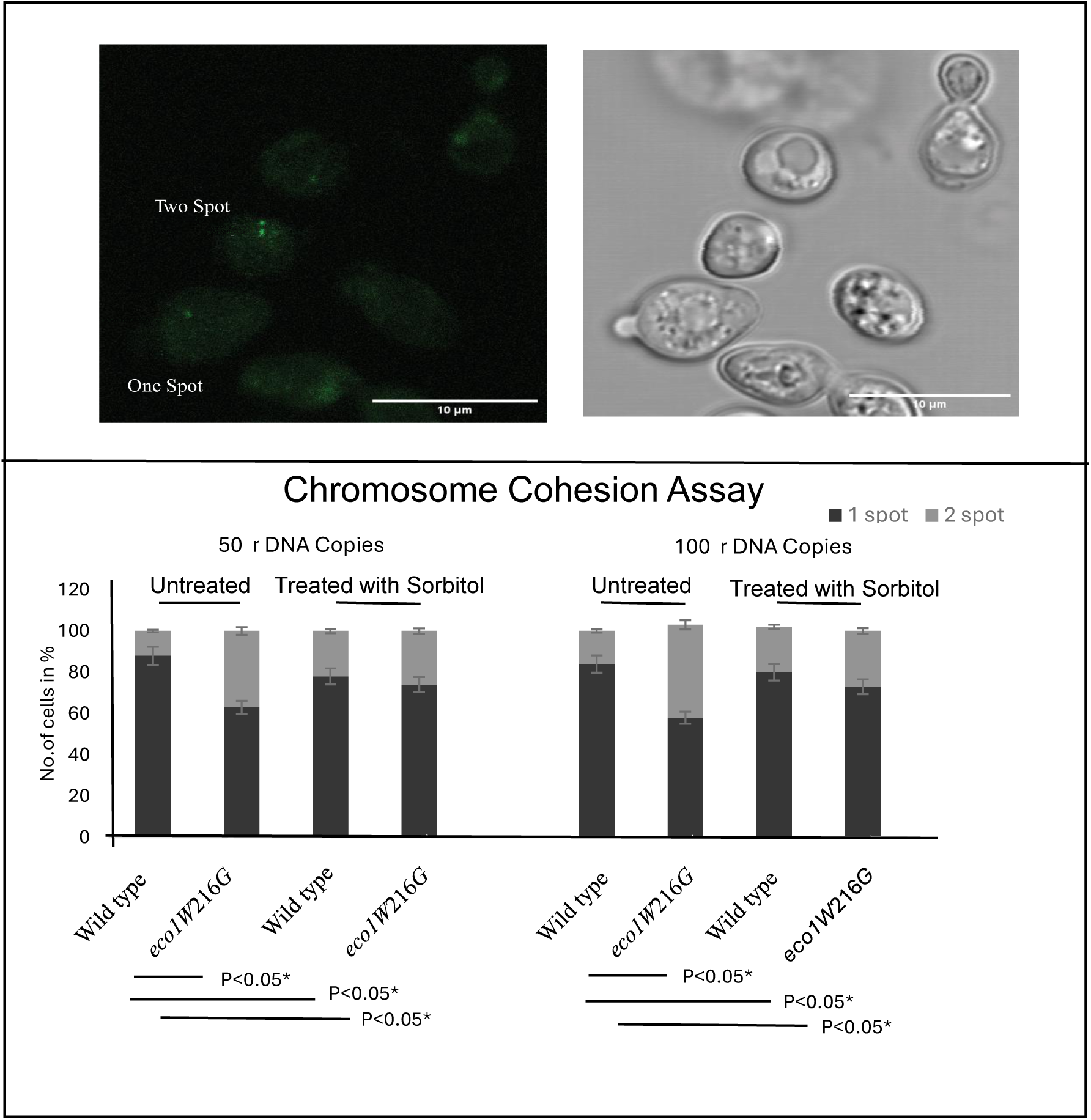
One Spot Two Spot Assay to check the effect of sorbitol on chromosome cohesion. Cohesinopathy mutants show defects in chromosome segregation. Wild type and mutant strains of *Saccharomyces cerevisiae* with 50 and 100 copies of rDNA were grown in YPD medium with and without 1.5M sorbitol at 30°C. The cells were nocodazole treated, to arrest them at metaphase and checked for the presence of one spot vs two spots. Images for live cells were collected using confocal microscopy with 100x objective. In order to quantify fluorescence intensity, approximately 100 cells of each type were counted. Graphs were prepared using GraphPad prism 9 software. T-test was used to calculate P values. (p <0.05*).

### 3.3. Effect of sorbitol on stress response genes

We wanted to check if there are changes in expression levels of the ER stress response genes in wild type and mutant, treated with sorbitol. We used candidate genes der1, kar2 and ero1. RT-PCR was done in different sorbitol concentrations of wild type and *eco1W*216*G*. The expression level of these target genes, showed that, in comparison to sorbitol-treated samples, there is little to no change in gene expression levels of wild type. In mutants, the increased expression of der1, ero1 and kar2 genes are corrected when treated with sorbitol **(Fig 3a, Suppl. Fig 1)**.

**Fig.3(a).**
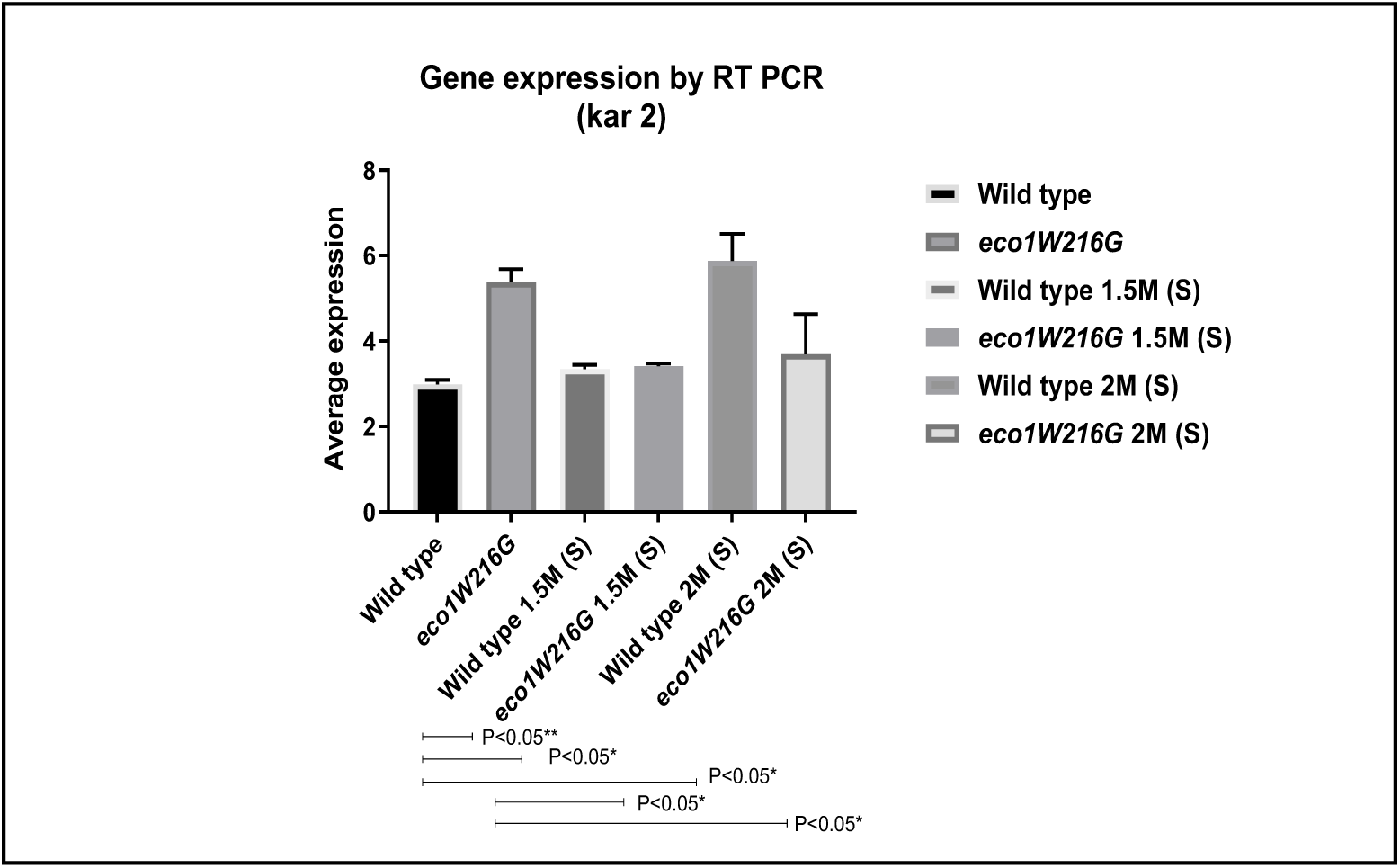
Effect of sorbitol on stress response gene. Data denoting fold change expression levels of stress response gene (kar2). RT-PCR was done with RNA from treated and untreated wild type and mutant cells. cDNA was prepared from the RNA extracted from wild type and mutant cells. Graphs were plotted from data obtained using GraphPad Prism 9 software. P values were calculated, all p-values are statistically significant. (P value<0.05*), act1 was used as control.

To analyze the overall changes in gene expression in presence of sorbitol, we performed RNA sequencing. In the gene expression study, the data showed that overall 541 genes were upregulated and 291 genes downregulated. To identify the genes whose expression levels were changed, we compared upregulated and downregulated genes with p<0.05 **(Fig 3b).** Out of the genes upregulated many of them belong to the ER stress response pathway and the others were ribosomal genes ^4^. In mutants treated with sorbitol, these were corrected to a certain level.

**Fig.3(b).**
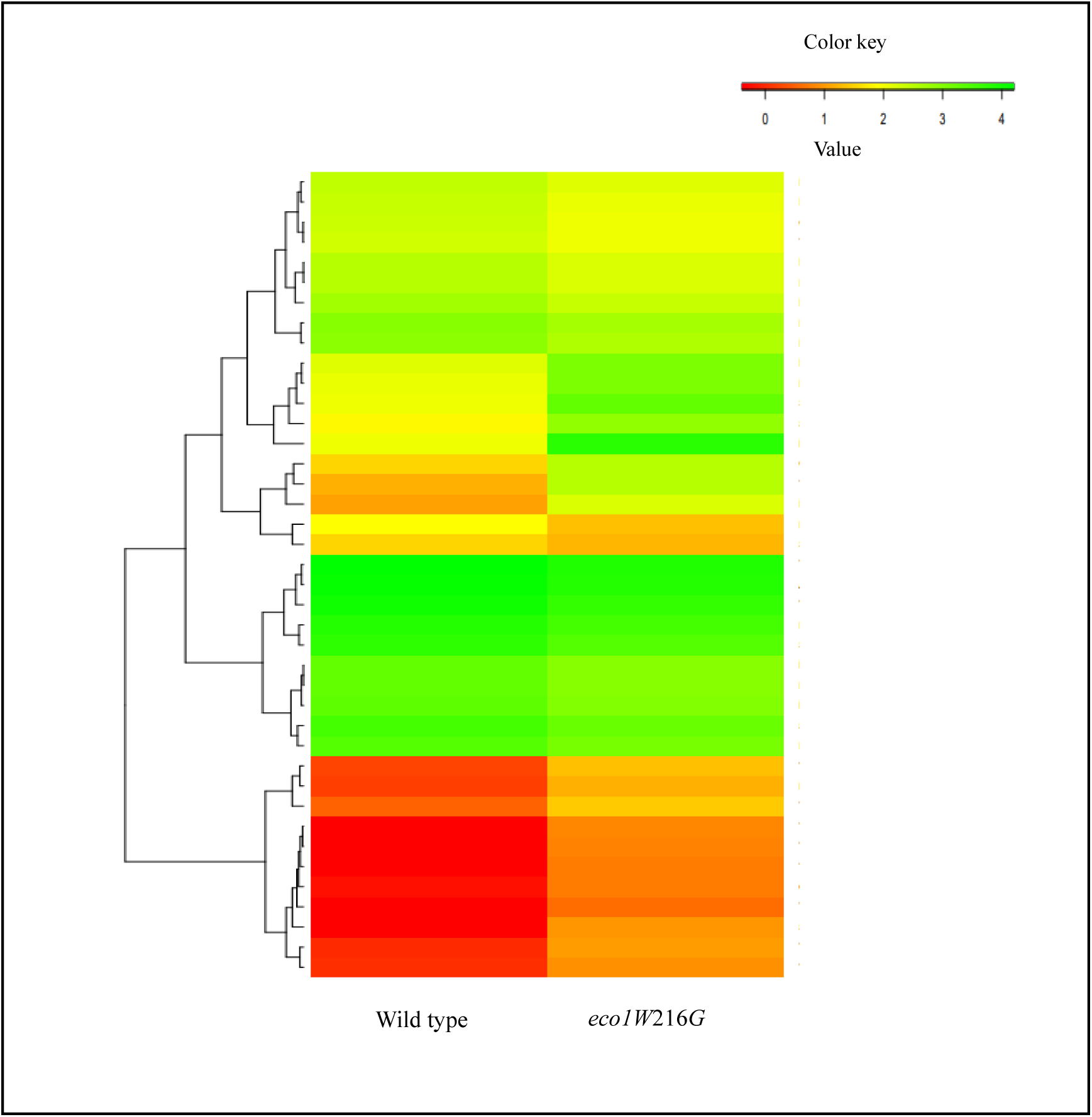
Heatmap showing gene expression in presence of sorbitol. Heatmap showing RNA sequencing based on gene expression pattern. Yellow and green values represent up and downregulated genes respectively. Color density indicates levels of fold change.

### 3.4. Luciferase sensitivity assay to detect if sorbitol can correct misfolding of proteins and reduce proteostasis stress

Chemical chaperones play an important role in proteostasis^49^. The defects associated with proteostasis is observed in many neurodegenerative diseases. To measure the proteostatic effects, we have used sensor protein from firefly^49^. We have used GFP-tagged protein constructs to detect protein misfolding, followed by protein aggregation. Slight change or imbalance in proteostasis changes the solubility of the luciferase reporter. This is detected by luciferase sensitivity assay **(Fig 4)** which shows that the luciferase activity in mutants was lower in comparison to wild types. On treatment with sorbitol, the luciferase activity increased in the eco1 mutant and almost reached wild type levels of the treated samples.

**Fig.4.**
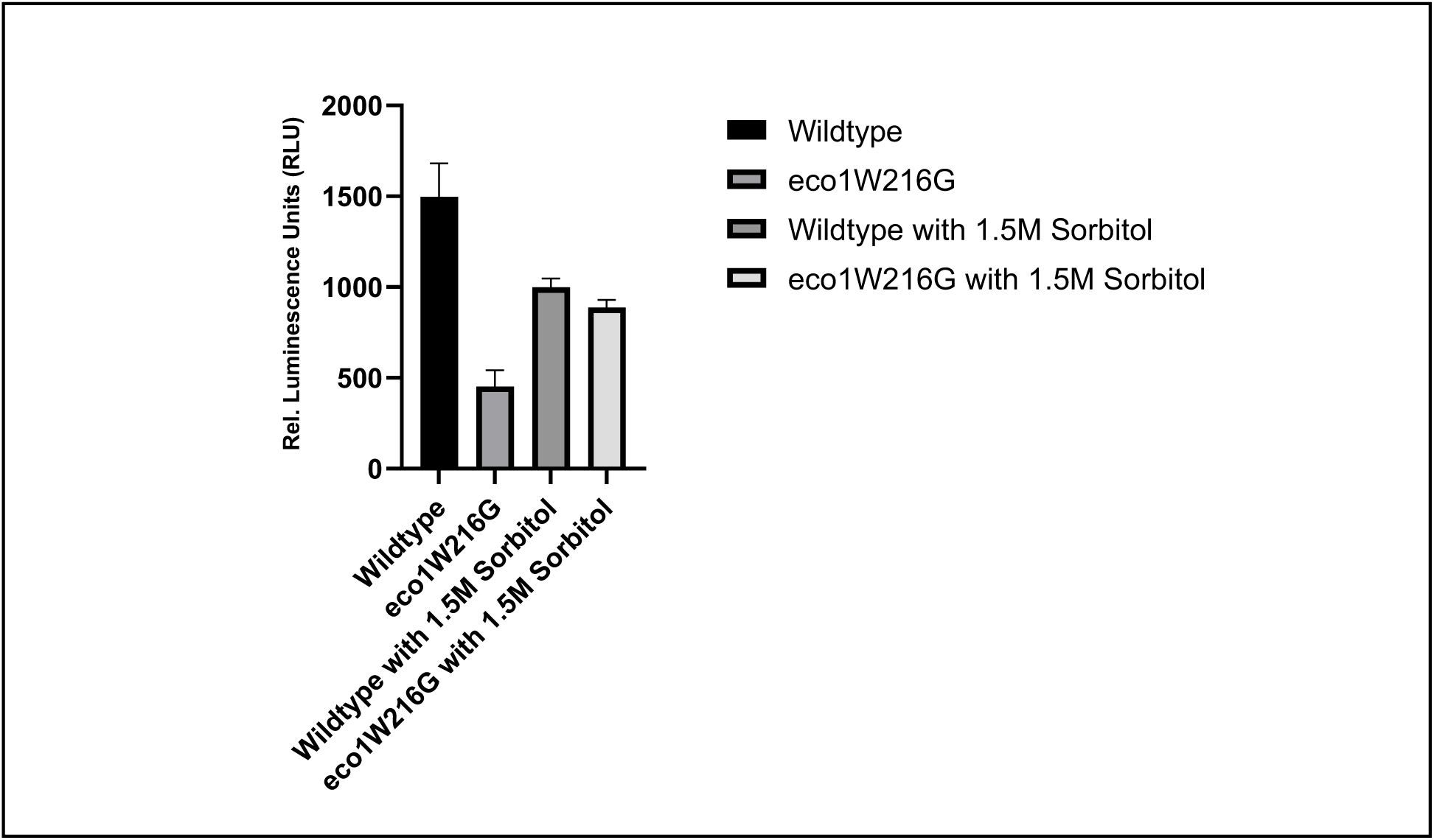
Luciferase reporter assay to check levels of proteostatic stress. The graph showing levels of proteostatic stress of wild type and mutant in presence and absence of sorbitol. The level of luminescence measured in luciferase assay indicates fold change of luciferase activity.

### 3.5. Changes in actively translating ribosome fraction in the presence of sorbitol

Polysome profiling is a technique that detects the association of mRNA with that of ribosomes and gives an idea about the overall translation taking place in the cells ^44^. This can also be used to detect the step of translation that gets affected ^50^. In case of defects in translation initiation polysome to monosome ratio decreases. On the contrary, if there is a translation elongation defect, the polysome to monosome ratio increases.

Here, we observed that peaks for polysomes are more pronounced in the wild type than in the mutant **(Fig 5)**. It is already known that polysome profile changes during stressed conditions ^46^. We have performed polysome profiling of wild type and mutant cells treated with 1.5M sorbitol and compared them with that of untreated wild type and mutant samples. After calculating the polysome to monosome ratio we observed that the polysome to monosome ratio of the mutants treated with sorbitol is similar to the values of wild types, which indicates that addition of sorbitol, helps in rescuing the mutant from translational impairment.

**Fig.5.**
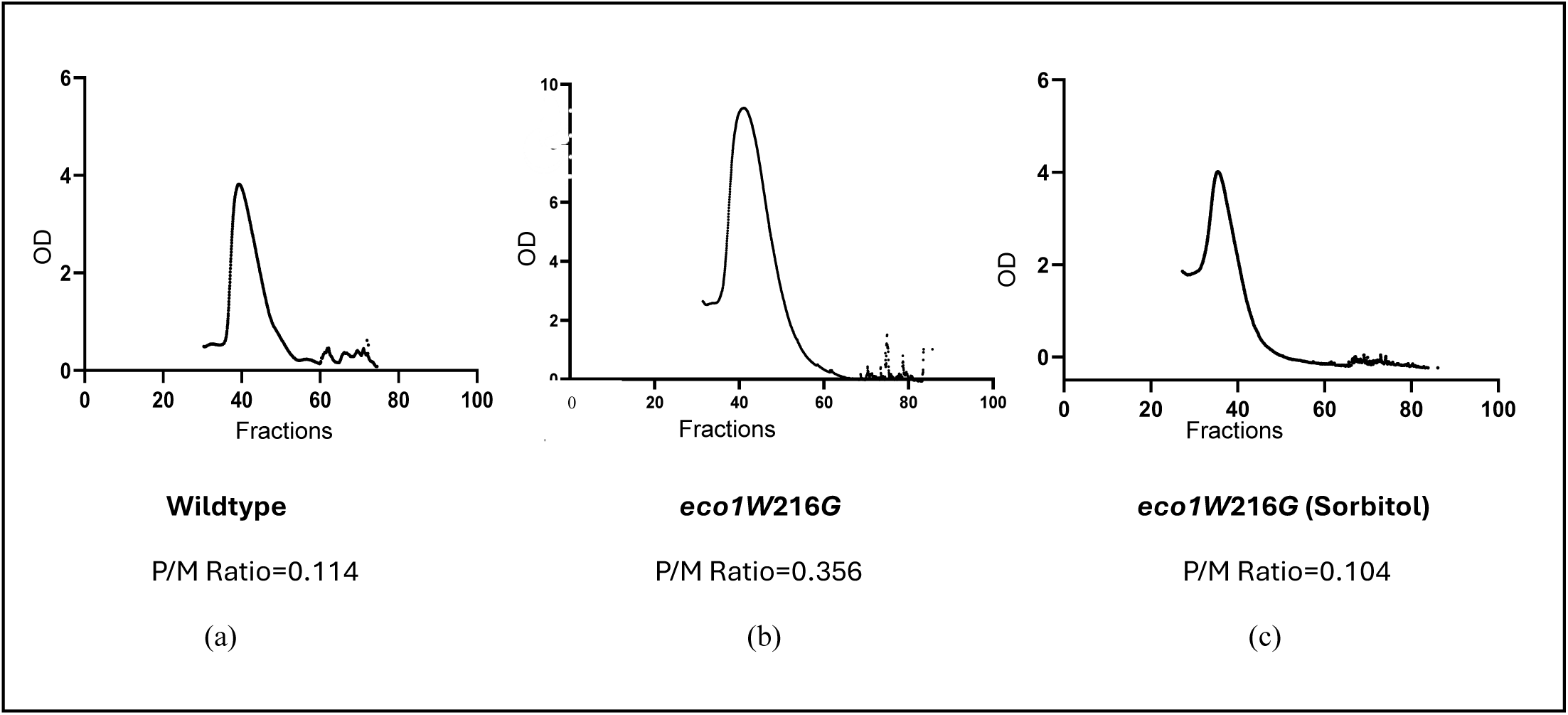
Effect of sorbitol on protein translation using polysome profiling. Polysome profiles of wild type and mutant strains grown in YPD, with and without 1.5M sorbitol at 30°C. Ratio of polysomes (P) to monosome (M), (P/M) is shown. **Fig5a.** Polysome profile of wild type, **Fig5b.** *eco1W*216*G* **Fig5c.** *eco1W216G* treated with 1.5M sorbitol.

### 3.6. Does sorbitol correct impaired protein translation in Cohesinopathy mutant strains?

Phosphorylation of eif2α is an indication of inhibition of translational initiation and is conserved during the process of evolution. eif2α phosphorylation of the ternary complex, leads to an inhibition of exchange from GDP to GTP, hence translation gets stalled ^4^. Here we have used immunoblotting techniques to measure the levels of phosphorylation of eif2α with eif2α alone as the loading control **(Fig 6)**. Analysis of the western blot image shows increased levels of phosphorylation of eif2α in the mutants **(Fig 6)**, is corrected in the samples treated with sorbitol.

**Fig.6.**
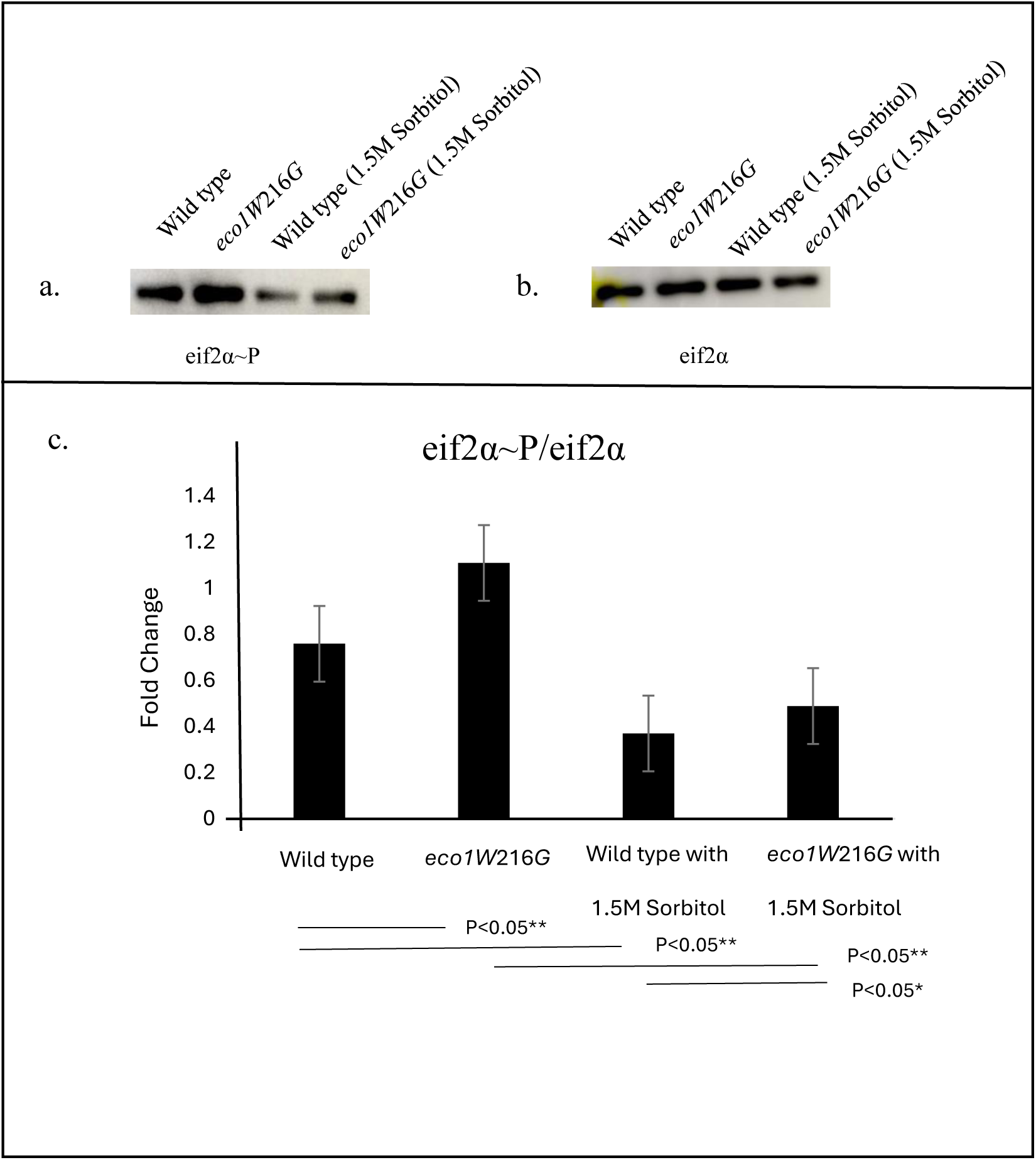
Effect of sorbitol on protein translation using western blot analysis. Whole cell extracts of wild type and *eco1W216G* grown in YPD broth with and without sorbitol. Protein extracts were used for western blotting to measure levels of eif2α and phosporylated eif2α, which is an indicator of translational inhibition. **Fig6a.** Western blot showing levels of phosphorylation of eif2α where wild type and *eco1W216G* untreated samples were used as control and wildtype and *eco1W*216*G* cells were treated with 1.5M sorbitol respectively. **Fig6b.** Expression level of eif2α (loading control)**. Fig6c.** Graph showing densitometry data.

### 3.7. Sorbitol might play a role in regulating autophagy

Autophagy is a recycling process of the cell which helps the cell to survive under stressed conditions. During stressful conditions, cells are deprived of nutrients. By the mechanism of autophagy, the breakdown and reuse of various proteins and macromolecules take place in the cells. The Roberts’ mutants are associated with translational defects, which probably affect translation initiation and ER stress. This might result in the accumulation of a higher amount of misfolded proteins ^13^. This increase in the levels of misfolded proteins is harmful to the cell. Therefore, cells use autophagy mechanisms, to eliminate misfolded and aggregated forms of proteins. Here we hypothesize, that the use of the chemical chaperone sorbitol may affect levels of autophagy in the mutant strains. In untreated samples, the levels of autophagy in the mutants are lower than in wild type strains by 40%. We have checked levels of ATG8 to determine if autophagy increases or decreases in presence of sorbitol. ATG8 is found in all membranes during autophagosome formation. Hence it is widely used to determine levels of autophagy. Our data suggests, that there is an increase in the levels of autophagy in the mutant strain treated with sorbitol than in the untreated cells, whereas, in wild types, the levels are almost similar **(Fig 7)**. We have used GAPDH **(Fig 7)** as the loading control.

**Fig.7.**
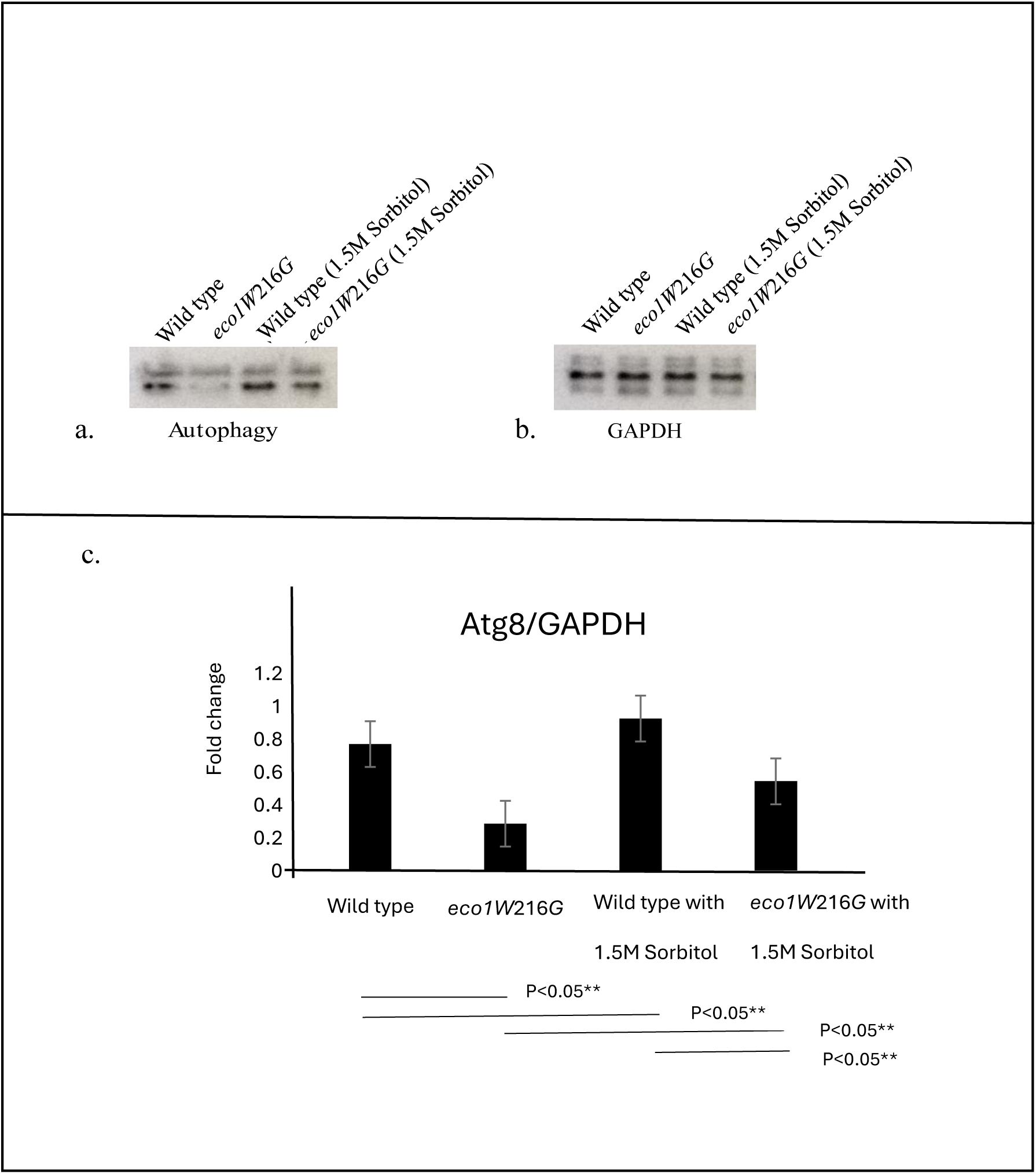
Effect of sorbitol on protein translation and autophagy using western blot analysis. Whole cell extracts of wild type and mutant cells were made from cells grown in YPD broth with and without 1.5M sorbitol. Lysates were used for western blotting to measure levels of autophagy using ATG8-GFP as a marker. Fig 7a. Western blot showing levels of ATG8-GFP with wild type (W303a) and mutant (*eco1W*216*G*) as control and wild type (W303a) vs mutant (*eco1W*216*G*) cells treated with 1.5M sorbitol. Fig 7b. Western blot of loading control using GAPDH. Fig 7c. Graph showing densitometry data.

**Fig. 8.**
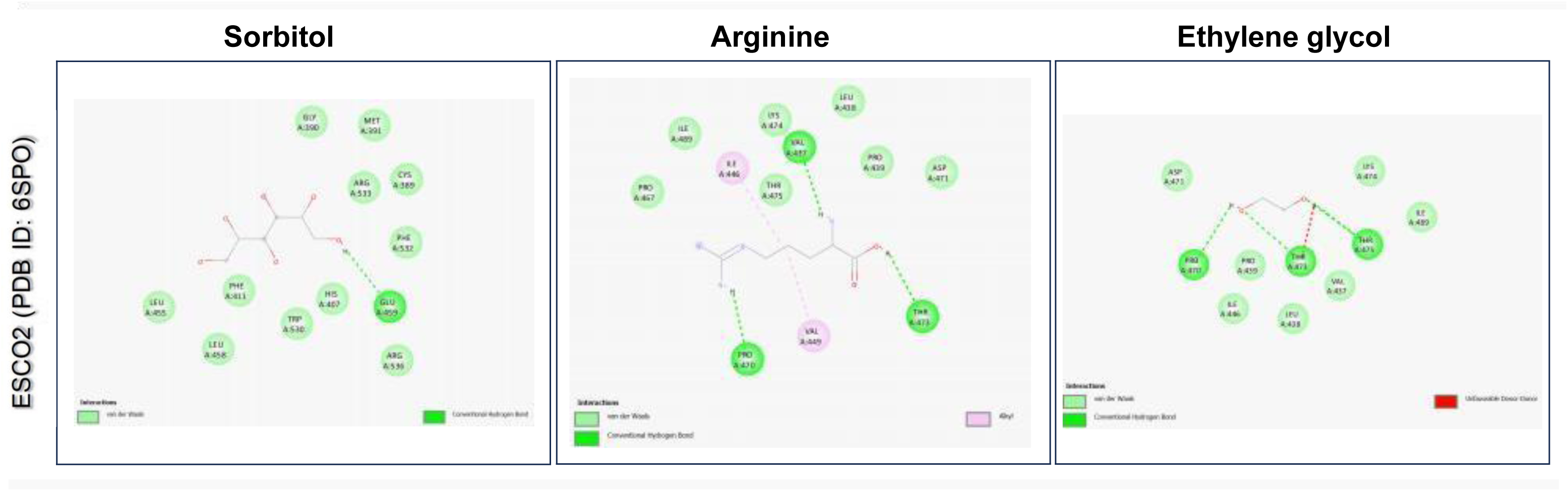
2D interactions of chemical chaperone sorbitol, arginine and ethylene glycol with ESCO2 (PDB ID: 6SP0).

To summarize the data for growth assays, it can be hypothesized, that the Roberts’ mutant is lethal at 37°C, but in presence of 1.5M sorbitol, rescues the mutant **(Fig 1).** In the chromosome cohesion (one spot two spots) assay, we find a reduction in the number of two spots after treatment with sorbitol **(Fig 1),** which indicates that chromosome cohesion is restored.

Results from RNA sequencing and RT-PCR showed reduced expression of stress response genes **(Fig 3)**. ER stress response gene levels were initially higher in the mutant strains than in wild type. Post-treatment with 1.5M and 2M sorbitol, the level of expression of the stress response genes is reduced in the mutant strain compared to untreated samples. After treatment with sorbitol, luciferase activity increases indicating a reduction in proteostastic stress **(Fig 4).**

To analyze the effect of sorbitol on translation, polysome profiling was done using a sucrose density gradient. The polysome to monosome ratio returns to almost wild type levels in sorbitol-treated samples with pronounced polysomes **(Fig 5)**. Taking cues from here, we wanted to determine the levels of eif2α phosphorylation. It was previously reported that eif2α phosphorylation increases in the *eco1W216G* mutant. This is in accordance with earlier literature which cites that a decrease in the polysome to monosome ratio is an indication of impairment of translation initiation. Translation initiation being affected through phosphorylation of eif2α, we found, in the sorbitol-treated mutants, the levels of phosphorylation of eif2α were reduced by 75% **(Fig 6)** in comparison to the mutant.

Proteostasis involves two important interdependent processes, one is phosphorylation of eif2α and the other is autophagy. The cells control the process of protein synthesis and autophagy by regulating the phosphorylation of eif2α^51^. Here cells restrict the rate of protein synthesis by phosphorylation of eif2α and improve energy utilization through autophagy. As a result, the load on protein folding and protein degradation machinery are diminished.

Our studies indicate that phosphorylation of eif2α and changes in translation can be correlated with autophagy. To understand the regulation of autophagy better, we have used immunoblotting techniques for checking the levels of ATG8 (Mammalian homologue-LC3, GABARAP, GATE-16 and ATGL8). The levels of ATG8 **(Fig 7)** are reduced in the Roberts’ mutant but increases to wild type levels in sorbitol-treated samples. It is found that levels of phosphorylation of eif2α in mutant strain is reduced, in the presence of sorbitol. On the contrary ATG8 levels are increased in the mutants when treated with sorbitol, similar to wild type conditions. The inverse correlation of eif2α and ATG8 levels give us an indication of the link between the alteration in translation machinery and autophagy, in sorbitol-treated samples. These pathways are possibly helping the cell in stressed conditions, when treated with sorbitol.

### 3.8 *In-silico* screening

To further understand the interaction between ESCO2 protein and sorbitol, molecular docking was conducted with sorbitol as a ligand, targeting esco2 in complex with Coenzyme A (PDB ID: 6SP0). Additionally, two known chemical chaperones—Arginine (as a positive standard) and Ethylene glycol (as a negative standard)— were included for comparison. Among the three chaperones, Arginine exhibited the highest binding affinity at −5.3 kcal/mol, while sorbitol showed a binding affinity of −4 kcal/mol (Table 1). Ethylene glycol displayed the lowest binding affibity of −3.3 kcal/mol. The binding interactions of the three chaperones demonstrated stability through hydrogen bonds with various amino acid residues of enzymes (**Table 1., Fig 8**). Arginine and ethylene glycol showed similar amino acid residues for hydrogen bonding (P470 and T473), whereas sorbitol specifically bonded with glutamic acid at position 459 (E459).

**Table 1.** Binding affinity of sorbitol with ESCO2 in complex with Coenzyme A (CoA) (PDB ID- 6SP0) and its interactions (PDB ID 6SP0) and its interactions.

| Compounds | Binding Affinity $\Delta G$ (in kcal/mole) with acetyltransferase in complex with CoA(PDB ID- 6SP0) | Interactions with acetyltransferase | |
| --- | --- | --- | --- |
|  |  | Other interactions | Hydrogen Bonding |
| Sorbitol | - 4.0 | Cys389, Gly390, Met391, Phe411, His437, Leu455, Leu458, Arg533, Phe532, Trp530, Arg536 | Glu459 |
| Arginine | -5.3 | **Leu438, Pro439,** Pro467, **Asp471,** Lys474, Thr475, Val449, **Ile489** | **Pro470,** Val437, Thr473 |
| Ethylene Glycol | -3.3 | **Asp471,** Lys474, **Ile489, Pro439,** Ile446, **Leu438,** Val437 | **Pro470,** Thr475, **Thr473** |

## 4. Discussion

The chaperones that are naturally present in cells, play a very important role by helping proteins to fold and maintain their native conformation. It also helps to remove misfolded proteins by activating the UPR mechanism ^52^. Sometimes, in presence of excess misfolded proteins, it increases the load of the chaperones and the protein degradation machinery. This results in ER and proteostatic stress ^23^. To reduce ER stress, we can use different chemical chaperones that can be added externally. Work on sorbitol by other groups suggests, that, it increases the solubility of cellular aggregates and facilitates the ER to Golgi transport ^36^. It also allows proper folding of proteins. Studies have been reported where sorbitol and trehalose reduce fibrillation in proteins ^53^. Bioinformatics analysis on the interaction of amyloid fibrils with sorbitol suggests changes in peptide conformation, kinetics and physical characteristics of the protein ^53,54^. Hence, we have selected sorbitol as the chemical chaperone.

In this report, we have used spot dilution and growth curve assay to check the effect of sorbitol on the growth patterns of wild type and *eco1W*216*G* strains at permissive and nonpermissive temperatures. Interestingly, screening these temperature-sensitive mutants in the presence of sorbitol, we found that the mutant strain rescues temperature sensitivity at 37°C in presence of sorbitol **(Fig 1)**. We have checked the growth phenotype of the eco1 mutant at different concentrations of sorbitol, upto 2M concentration. At this concentration, it shows better rescue of the mutants. Simultaneously, it is observed that the growth of wild type cells is also affected to a certain extent. Hence, we have selected 1.5M sorbitol concentration for further studies.

Similar rescue in temperature sensitivity has been observed by other groups working on the cohesin mutant *mcd1-1* mutant ^2^, treated with sorbitol. They have suggested that the rescue, could be due to correction of cohesion defects ^2^. To check if the addition of sorbitol helps in rescuing cohesion defects we have performed (one spot two spot) chromosome cohesion assay ^4^. The data obtained from this assay shows that the number of two spots are reduced in samples that are treated with 1.5M sorbitol (**Fig 2**). Since the presence of two spots indicates, precocious separation of sister chromatids, we may conclude that, addition of sorbitol rescues the cohesion defects in the *eco1W*216*G* mutants.

Every cell responds to environmental stress by altering the expression levels of certain proteins that are required by the cells to adapt to stressed situations. This, in turn, helps the survival of the cell ^53^. Usually, this alteration in protein expression is controlled by expression levels of mRNA and altering attachment of mRNA to ribosomes ^54^. Cells under favorable conditions can grow optimally, but when they are under stress, some stress response genes get upregulated ^55^. In this study we have used sorbitol, to find if there are changes in the expression pattern of the genes. We have also tried validating the levels of the stress response gene in presence of sorbitol by RT-PCR. The results indicate that the expression levels of kar2, der1 and ero1, ER stress response genes, elevated in the mutant, are reduced in presence of sorbitol **(Fig 3)**.

In certain cases, during stressed situations, various degradation pathways like proteosomal and autophagic mechanisms, are upregulated, whereas pathways that control protein synthesis are downregulated ^16^. The effect of sorbitol on proteostatic stress was measured by luciferase assay and results showed increased luminescence in mutant samples, indicating reduction in proteostatic stress **(Fig 4)**. We have subjected the wild type and mutant cells to different concentrations of sorbitol and checked their effects on growth, transcription, translation, protein folding and its effect on autophagy.

The eco1 mutant generates ER stress and in response to this stress, there is a change in the translation pattern ^56^. It was important to check the polysome profile of these cells. Earlier reports showed that the polysome profile in the eco1 mutant are different in comparison to that of the wild type. The polysome to monosome ratio is reduced in the mutant. When treated with 1.5M sorbitol, polysome to monosome ratio tends to be similar to that of the wild type **(Fig 5)**.

Protein synthesis is controlled at various steps ^57^. During translation initiation, phosphorylation of translation initiation factor eif2α, is one of the key controlling steps. Phosphorylation of eif2α is associated with reduced protein synthesis ^58^. Thus it is considered as an indicator of translation inhibition which is evolutionary conserved ^4^. Our observations from the western blot suggest, that in presence of 1.5 M sorbitol, the levels of phosphorylation in the cells decrease in the mutant when treated with sorbitol compared to the control strains. In presence of sorbitol, eif2α phosphorylation is inhibited. Data from **Fig 6**, shows phosphorylation levels were elevated by 75%.

Autophagy is the pathway known for the degradation and reuse of proteins playing important roles during various cellular processes such as nutrient starvation and misfolded protein accumulation ^23,24,27^. It has been observed that during hyperosmotic stress, genes for LC3 (Autophagosomal ortholog of yeast ATG8) ^59^ get induced and trigger autophagy-related protein (ATG8) production to clear protein aggregates and rescue the cells from a stressed situation ^60^. We wanted to check, if there is an induction of autophagy, in the presence of sorbitol. Our data suggests, that levels of autophagy are higher in mutants treated with sorbitol than in untreated samples **(Fig 7)**. At present ATG8 is the most widely used organelle marker to determine autophagy ^61^.

In contrast to the results of eif2α phosphorylation, we find levels of ATG8 increases in the mutant treated with sorbitol. In a separate study, in nutrient-starved yeasts, it was found, that phosphorylation of eif2α is elevated, whereas autophagy detected from expression levels of ATG8 is reduced ^62–64^. This data supports our findings. In the *eco1* mutant, we find eif2α phosphorylation increases and autophagy decreases as evidenced by levels of ATG8. In cells treated with sorbitol, we find that the levels of eif2α phosphorylation are reduced and autophagy is restored to wild type conditions. The increased levels of autophagy is an indication of excess ER stress or external osmotic pressure built up in the cells ^64^. Under normal conditions, autophagy increases to counteract the proteostatic stress, leading to clearance of accumulated protein aggregates. However, in the eco1 mutant, autophagy being suppressed, the proteostatic stress level remains elevated.

Like trehalose, there is a possibility, that the presence of sorbitol may play a chaperone-like function to lower ER stress, which possibly initiates autophagy in cells ^65^.

To correlate the activity between eif2α phosphorylation and autophagy, we found increased levels of eif2α phosphorylation ^56^ and a decrease in autophagy as evidenced by expression levels of ATG8 ^66^. When these mutants, were treated with sorbitol, we found a decrease in the phosphorylation levels of eif2α and ATG8 levels were enhanced ^51^. The polysome profile reverted to wild type conditions in the presence of sorbitol.

*In silico* studies of sorbitol, with eco1 (human esco2), helps us predict possible binding interactions. ESCO1 and ESCO2 proteins belongs to the GNAT (GCN5-related N acetyltransferase) family with N terminal C_2_H_5_ zinc finger and C-terminal acetyltransferase domain (**Fig 8**). ESCO1 protein is constantly expressed whereas ESCO2 protein levels are high during S-phase of cell cycle ^67^. These proteins stabilize cohesin proteins on DNA that help to align the sister chromatids during mitosis. Cohesin is a complex ring like structure made up of SMC (structural maintenance of chromosomes) proteins comprising Smc1, Smc3, Scc1 (Mcd1) (kleisin), and SA1/SA2. Acetylation of basic patch of Smc3 domain by ESCO proteins weaken DNA binding of release factors, which is stabilized by Wapl-Pds5 accessory factors, thus stabilizing cohesion^68–70^. So, ESCO proteins, particularly ESCO2, is crucial for segregation of sister chromatids. There are in total 26 known mutation of ESCO2 proteins that leads to complete or partial loss of acetyltransferase activity ^5,71^. The ESCO2 mutant (W539G) results in loss of autoacetyltransferase activity ^72^. Molecular docking findings demonstrated, sorbitol interacts with acetyltransferase domains of protein residues, including R533, F532, W530, R536 environment near amino acid W539 unlike arginine or ethylene glycol. These interactions with sorbitol might aid in restoring acetyltransferase activity and rescue the temperature sensitivity. The molecular dynamic investigation might give further insight for these interactions and restoration of acetyltransferase activity.

Summarizing these results, we can conclude, that use of sorbitol rescues the Roberts’ mutant from stressed condition and corrects the global protein translation, misfolding and autophagy along with chromosome cohesion defects (**Fig9**).

**Fig. 9.**
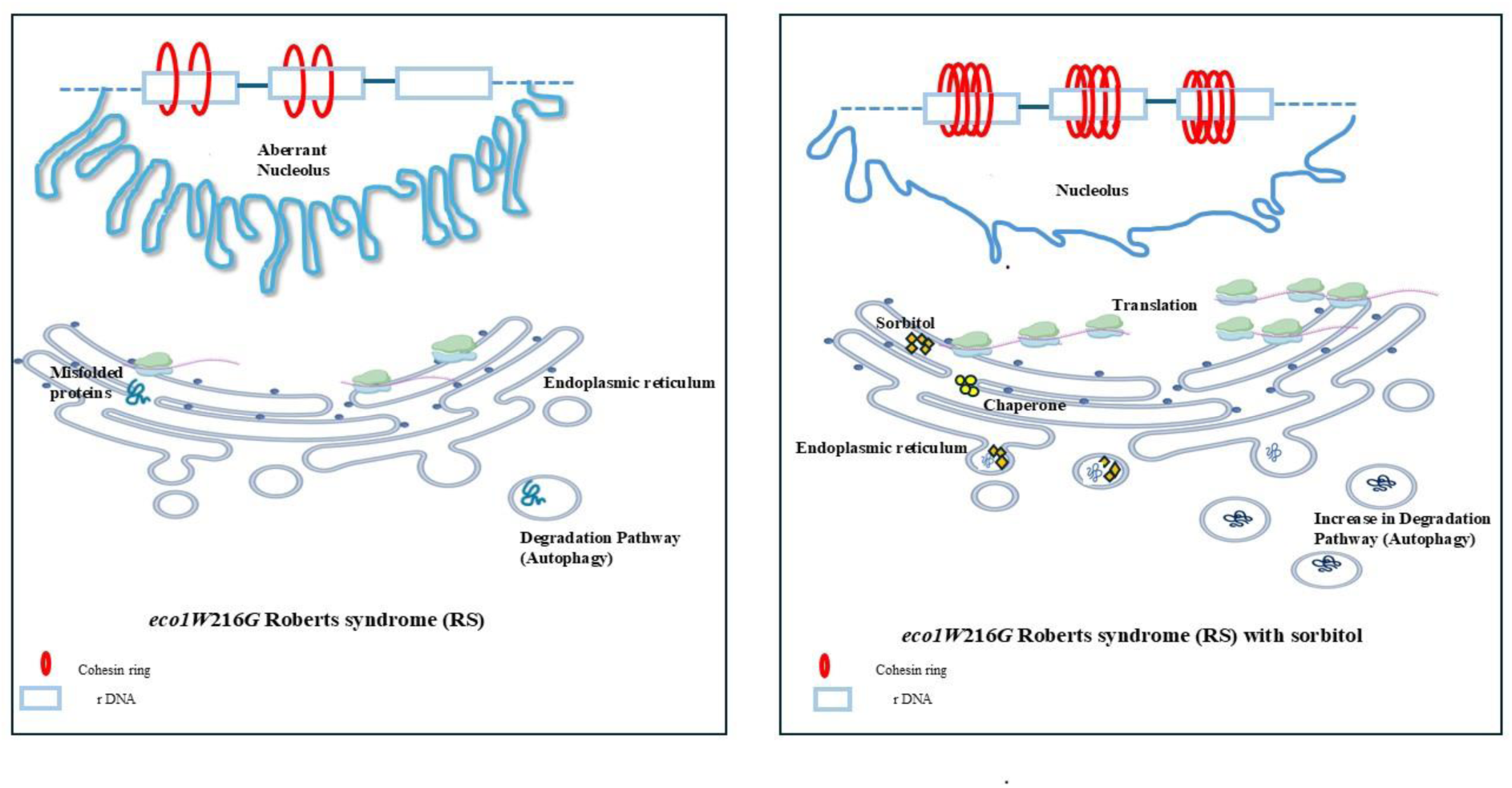
Schematic of Hypothetical Model. (a) *eco1W*216*G* shows disrupted nucleolus and reduced cohesion at rDNA, which in turn impairs translation, protein misfolding and autophagy. (b) Roberts mutant treated with sorbitol shows conspicuous nucleolus with increased cohesion at rDNA, correction of translation, protein misfolding and enhanced autophagy.

## 5. Conclusions

From our work, we can hypothesize that sorbitol is helping regulate chromosome cohesion, translation, protein folding and autophagy associated errors This, in turn allows proper chromosome segregation in the sorbitol-treated Roberts’ mutant. Reduced levels of autophagy in mutant cells cause excess protein built up ^73^, but the addition of sorbitol leads to an increase in levels of autophagy which in turn might reduces the protein accumulation load. ^74^. Thus, we hypothesize that sorbitol acts as a chemical chaperone which might interact with the protein molecule and help it fold to maintain its native structure. This stabilizes proteins and prevents them from forming aggregates as well as corrects other downstream effects.

## Data Availability Statement

Strains are available on request. The authors affirm that all data necessary for the conclusions of the article are present in this article, figure and supplementary material.The raw RNA Sequencing files are deposited in the SRA (Sequence Read Archive) with BioProject database (https://www.ncbi.nlm.nih.gov/bioproject/PRJNA1151841).

## Acknowledgements

We would like to thank Dr. Jennifer Gerton for the *eco1W*216*G* strain.

## Author Contribution Statement

The experiments were designed by TB and AM. Experiments and analysis were performed by SM, TB, AM, VN and HC. MS was prepared by TB, AM, HC, VN and SM.

## Abbreviations Used

WT: Wild type
RS: Roberts syndrome
ER: Endoplasmic reticulum
UPR: Unfolded protein response
CWI: Cell wall integrity
eif2α: Eukaryotic initiation factor 2

