## Supplementary material for "Polyol sugar osmolyte - sorbitol corrects chromosome cohesion and translational defects in cohesin mutants": supply file

**Supplymental data**

**Figure S1**

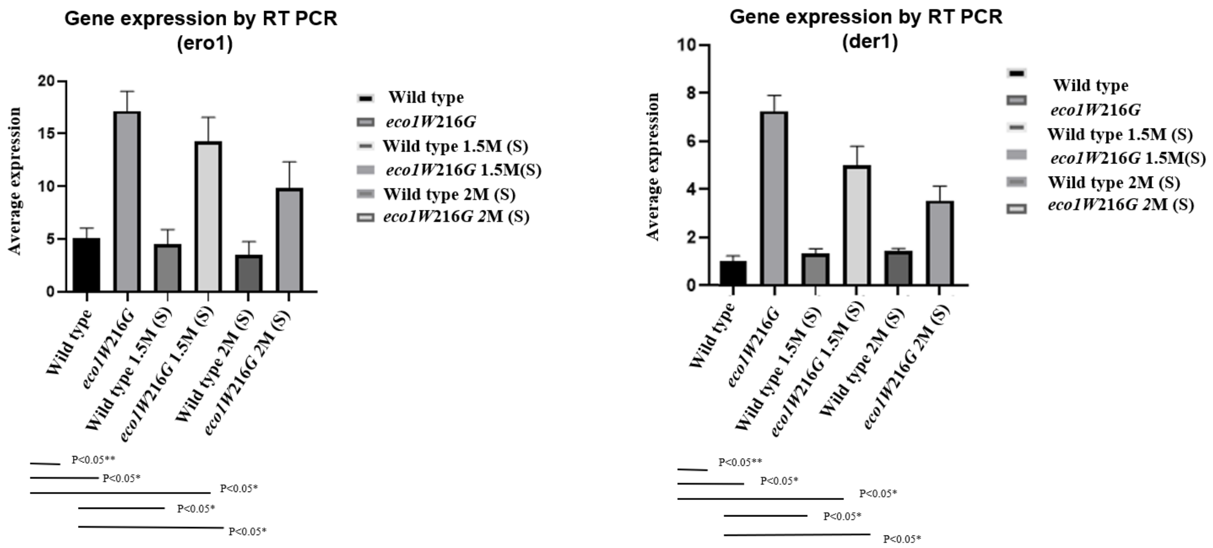


**Effect of sorbitol on stress response gene**

Data denoting fold change expression levels of stress response gene (ero1,der1). RT-PCR was performed on RNA from treated and untreated wild type and mutant cells. cDNA was prepared from RNA extracted from wild type and mutant cells grown at 30°C temperature in YPD medium with and without 1.5M and 2M sorbitol. Graphs were plotted from data obtained using GraphPad Prism 9 software. P values were calculated, all p-values are statistically significant. (P value<0.05*), act1 was used as control.

**Table S1**

| Figure | Strain Name | Genotype |
|---|---|---|
| Wild type | Wild type | MATa bar1-ura3 leu2 trp1 lys2 ade2 his3::pCUP1- GFP12-LacI12::HIS3 arm IV::LacO-URA3 |
| eco1W216G | eco1W216G | MATa bar1-ura3 leu2 trp1 lys2 ade2 his3::pCUP1-GFP12- LacI12::HIS3 arm IV::LacO-URA3 ECO1::eco1-W216G::HYG |
| Fig.2 | YSI126 | YSI102 except his3-11::GFP-LacI-HIS3 RDN1::LacO(50)-ADE2 |
| Fig.2 | BH283.1 | YSI102 except his3-11::GFP-LacI-HIS3 RDN1::LacO(50)-ADE2 ECO1::eco1-W216G::HYG |
| Fig.2 | YSI129 | YSI105 except his3-11::GFP-LacI-HIS3 RDN1::LacO(50)-ADE2 |
| Fig.2 | BH284.1 | YSI105 except his3-11::GFP-LacI-HIS3 RDN1::LacO(50)-ADE2 ECO1::eco1-W216G::HYG |
